# Enhancing cGMP signaling with psilocybin reduces head twitch and restructures the synaptic proteome while maintaining antidepressant response

**DOI:** 10.64898/2026.03.06.710108

**Authors:** Gabriele Floris, Sarah J. Jefferson, Jocelyne Rondeau, Abigail L. Yu, Frank S. Menniti, Alex C. Kwan, Joao P. De Aquino, John H. Krystal, Christopher Pittenger, Alfred P. Kaye

**Affiliations:** Yale University Department of Psychiatry, New Haven, CT; Freedom Biosciences, Tiburon, CA 94920; George & Anne Ryan Institute for Neuroscience, University of Rhode Island, Kingston, RI; Meinig School of Biomedical Engineering, Cornell University, Ithaca, NY; Department of Psychiatry, Weill Cornell Medicine, New York, NY; Yale University Department of Psychology, New Haven, CT; Yale University Department of Neuroscience, New Haven, CT; Yale University Child Study Center, New Haven, CT; Yale University Center for Brain Mind Health, New Haven, CT; Yale University Department of Comparative Medicine, New Haven, CT; Yale University Wu Tsai Institute, New Haven, CT

**Author notes:** Equal Contributions (Gabriele Floris, Sarah J. Jefferson). These authors jointly supervised the work (Christopher Pittenger, Alfred P. Kaye). **Corresponding author:** Alfred P. Kaye.

## Abstract

Psilocybin has antidepressant effects, but its 5-HT_2_AR-mediated perceptual effects limit tolerability. We combined psilocybin with a phosphodiesterase-9 inhibitor (PDE9i) and observed suppression of head-twitch response, but maintenance of antidepressant-like behavior. Proteomics showed that PDE9i-psilocybin reduced 5-HT_2_AR-mediated pathways while enhancing synaptogenesis. These results suggest that PDE9i-psilocybin represents a promising therapeutic strategy.

## Introduction

Major Depressive Disorder affects over 300 million people worldwide and approximately one-third of them fail to respond to conventional antidepressants [1]. Psilocybin, a serotonin 2A receptor (5-HT_2_AR) agonist, shows marked efficacy in refractory depression in phase 3 clinical trials (NCT05711940) [2]. Psilocybin administration is accompanied by marked changes in perception mediated by 5-HT_2_AR-dependent disruption of functional brain networks, including the default mode network [3]. These acute psychedelic effects can limit broad clinical implementation, motivating efforts to dissociate therapeutic from psychedelic effects. Multiple strategies are being explored, including non-hallucinogenic analogs, biased agonists targeting specific 5-HT_2_AR signaling pathways, and structure-based drug design approaches [4-5]. Combination-based strategies offer an alternative approach that maintains psilocybin’s therapeutic efficacy, while pharmacologically targeting mechanisms underlying its acute perceptual effects. Phosphodiesterase-9A (PDE9A) represents a compelling molecular target in this context. PDE9A is a high-affinity, cGMP-specific enzyme (Km ≈ 70 nM) with dense expression in hippocampus and prefrontal cortex, where it regulates cGMP pools distinct from those controlled by other phosphodiesterases [6]. Selective PDE9 inhibitors, such as BAY 73-6691 and PF-04447943, enhance synaptic plasticity and produce robust pro-cognitive effects in rodent models [7]. PDE9 inhibition reverses 5-HT_2_AR-dependent mescaline-induced scratching behavior [8], providing direct evidence that cGMP modulation can prevent behavioral outputs of 5-HT_2_AR activation. Here, we investigated whether PDE9 inhibition modulates behavioral and molecular responses to psilocybin. We quantified head twitch response (HTR) as a behavioral readout of 5-HT_2_AR engagement and predictor of acute perceptual effects, and assessed whether PDE9 inhibition interferes with psilocybin’s antidepressant-like effects in a chronic stress paradigm. Finally, synaptic proteomic analysis was performed to characterize the molecular landscape underlying these combinatorial effects.

## Methods

### Animals and Housing

Male C57BL/6J mice (8–12 weeks old; The Jackson Laboratory) were group-housed under standard conditions (12:12 h light/dark cycle) with ad libitum food and water. All procedures were approved by the Yale University Institutional Animal Care and Use Committee.

### Drugs

Psilocybin (item #14041, Cayman Chemical Company) was dissolved in saline and administered at 1 mg/kg, i.p. The PDE9A inhibitor PF-4181366 was synthesized and generously provided by Yuelian Xu at Chinglu Pharmaceutical Research LLC. It was analyzed by UPLC-MS at 98.95% purity. PF-4181366 was dissolved in 5% DMSO, 5% Tween-80 in saline at doses previously shown to increase cGMP levels in cortex and hippocampus (0.32, 1, 3.2, or 10 mg/kg, i.p.) [9].

The psilocybin dose (1 mg/kg i.p.) produces robust 5-HT_2_AR-dependent HTR and neuroplastic responses in preclinical models [10]; human therapeutic doses span ∼0.14– 0.36 mg/kg orally with linear pharmacokinetics [11].

### Head Twitch Response

We detected HTR using automated magnetometer-based system. Psilocybin (1 mg/kg) was administered 30 min after PF-4181366 (0.32, 1, 3.2, or 10 mg/kg) or vehicle and HTR was quantified immediately thereafter.

### Locomotor Activity

Following 1h habituation, mice were placed individually into standard cages positioned within a 16-beam infrared photobeam grid (Med Associates Inc.). Locomotor activity was recorded for 60 min pre- and post-injection of PF-4181366 (0.32, 1, 3.2, or 10 mg/kg) or saline.

### Chronic Multimodal Stress

Chronic multimodal stress (CMMS) was adapted from Hesselgrave et al. [12]. Mice were single-housed and habituated to two-bottle choice. Baseline sucrose preference was assessed on Days 2–3 during 14h dark-phase sessions. Beginning Day 4, mice underwent 15 consecutive days of stress: 5h daily restraint in ventilated 50 mL conical tubes under concurrent 2 Hz strobe lighting and 75 dB white noise. Session timing varied daily (8-11 AM). Sucrose preference was reassessed on Days 18–19 (post-stress) and on Day 20 at 4h post-treatment.

### Sucrose Preference Test

Sucrose preference was calculated as [sucrose intake / (sucrose + water intake)] × 100. A 1.5% solution was used during habituation and 1% for testing. Bottle positions were alternated to control for side preference.

### Tail Suspension Test

On Day 17, mice were suspended by adhesive tape (1 cm from tail tip) to a horizontal metal bar. Behavior was video-recorded for 6 min, and immobility—defined as absence of active escape-oriented movements—was scored manually during minutes 1–5.

### Pharmacological Treatment

Following TST, mice were stratified by sucrose preference and immobility into three groups: (1) vehicle–saline, (2) vehicle–psilocybin (1 mg/kg), and (3) PF-4181366 (10 mg/kg)– psilocybin (1 mg/kg). A 30 min interval separated pretreatment from psilocybin injection.

### Forced Swim Test

On Day 21 (24 h post-treatment), mice were placed individually in cylindrical beakers filled with water (21 ± 1°C; 15 cm depth) for 6 min. Immobility—passive floating with minimal movements to maintain head above water—was scored during minutes 2–6.

### Synaptosome isolation and LC-MS/MS

Mice were euthanized and the mPFC including ACC, PL and IL were dissected on ice. Crude synaptosomes were isolated as described in [13]. LC-MS/MS data were acquired on a Thermo Scientific Orbitrap Fusion mass spectrometer coupled to a Waters M-Class UPLC system. Protein identification and label-free quantification (LFQ) were performed in Proteome Discoverer (v3.2, Thermo Fisher Scientific). Normalization was based on total peptide amount and abundance ratios on pairwise peptide ratios. Background-based t-tests were used to compare protein abundances between conditions. Peptide- and protein-level FDR was set to 1%; only proteins with >2 unique peptides present in >50% of samples were retained. Canonical pathway enrichment graphs and hierarchical clustering of pathways were generated through QIAGEN IPA (QIAGEN Inc., https://digitalinsights.qiagen.com/IPA). Kinase enrichment analysis was performed using KEA3 (https://maayanlab.cloud/kea3/) and protein association networks depicted using STRING (https://string-db.org).

### Analysis of single-cell transcriptomics data

Allen Institute SmartSeq single-cell RNA-seq data were analyzed as described in [14].

## Results

### Pharmacological inhibition of PDE9 attenuates psilocybin-induced head-twitch responses without affecting locomotor activity

Mice received psilocybin (1 mg/kg, i.p.) 30 min after vehicle or PDE9 inhibitor (PDE9i) PF-4181366 pretreatment (0.32, 1, 3.2, or 10 mg/kg), and HTR was recorded for 30 min. ANOVA revealed a significant main effect of pretreatment on psilocybin-induced HTR (F (4,23) = 6.042, p = 0.0018). PF-4181366 at 3.2 mg/kg and 10 mg/kg significantly reduced HTR relative to vehicle (p = 0.0097 and p = 0.0025, respectively), whereas lower doses produced nonsignificant trends (Tukey’s multiple comparisons test, both p = 0.0532). Expressed as percentage of vehicle control, PF-4181366 reduced psilocybin-induced HTR by approximately 47% at 0.32 and 1 mg/kg, 59% at 3.2 mg/kg, 69% at 10 mg/kg. Locomotor activity was unaffected by PF-4181366 at any dose tested (F(4,24) = 0.298, p = 0.877; all adjusted p ≥ 0.96; data not shown), indicating that HTR attenuation reflects specific modulation of 5-HT_2_AR-mediated behavior rather than nonspecific motor suppression.

### PDE9 inhibition does not interfere with the antidepressant-like effects of psilocybin in a chronic stress model

Antidepressant-like behavior was assessed 24 h post-treatment using the forced swim test (FST). ANOVA revealed a significant treatment effect on immobility (F (2,57) = 9.855, p = 0.0002). Psilocybin significantly reduced immobility relative to saline-treated controls (p = 0.0008). Critically, co-administration of PDE9i did not attenuate this effect—the combination group similarly exhibited reduced immobility compared with saline controls (p = 0.0004). Thus, psilocybin reduces depressive-like behavior following chronic stress, an effect maintained with PDE9i co-administration. Sucrose preference did not differ among vehicle, psilocybin, or psilocybin+PDE9i groups (two-way RM ANOVA: treatment, F(2,75) = 0.517, p = 0.599; time × treatment, F(3.26,122.3) = 0.405, p = 0.766; data not shown).

### Synaptosomal proteomics reveal that PDE9i–psilocybin combination enhances synaptogenesis-related signaling while reducing 5-HT_2_AR downstream pathways

Proteomic analysis of mPFC synaptosomes collected 24h post-treatment revealed that PDE9i–psilocybin significantly altered synaptic protein composition relative to psilocybin alone. Pathway analysis showed an overall pattern of pathway downregulation with PDE9i– psilocybin, including GPCR and serotonin receptor signaling. Remarkably, synaptogenesis, believed to be crucial for psychedelic therapeutic effects, was upregulated in the combination group. Kinase enrichment analysis predicted upstream kinases regulated by PDE9i-psilocybin. For upregulated proteins, top kinases were involved in synaptogenesis and structural remodeling including mTOR and Src. For downregulated proteins, top kinases included downstream targets of canonical Gq-coupled 5-HT_2_AR signaling such as PKC (Prkca and Prkcb). Taken together, these data suggest that PDE9i–psilocybin suppresses 5-HT_2_AR-mediated downstream signaling while preserving synaptogenesis. Analysis of Allen Institute’s single cell sequencing data showed co-expression of Htr2a and PDE9a transcripts in several classes of cortical glutamatergic neurons including PT neurons, which are required for psilocybin-induced structural plasticity [14].

## Discussion

These findings demonstrate that pharmacological inhibition of PDE9 attenuates the acute head-twitch response to psilocybin without compromising its antidepressant-like effects in a chronic stress model. This behavioral dissociation is mechanistically supported by synaptosomal proteomic analysis, showing preserved synaptogenic signaling pathways in the combination group, consistent with maintained therapeutic potential. The ability to reduce acute 5-HT_2_A receptor-mediated behavioral responses while preserving antidepressant efficacy suggests that PDE9 inhibition may represent a viable strategy for improving the clinical tolerability of psilocybin. Notably, the 24-hour post-treatment timepoint selected for proteomics aligns with the established onset of psilocybin-induced neuroplastic consolidation in humans, where neuroimaging studies report changes in brain network connectivity and clinical symptom relief emerging within 24 hours of dosing and persisting for at least one month—providing translational grounding for the molecular window captured here [15]. It is possible that the increase in cGMP following PDE9 inhibition functions to block psilocybin-induced hyperexcitability and thereby reduce head-twitch responses. Collectively, these results support the hypothesis that combination approaches targeting intracellular signaling can modulate the acute profile of psychedelics while preserving their therapeutic mechanisms.

**Figure 1.**
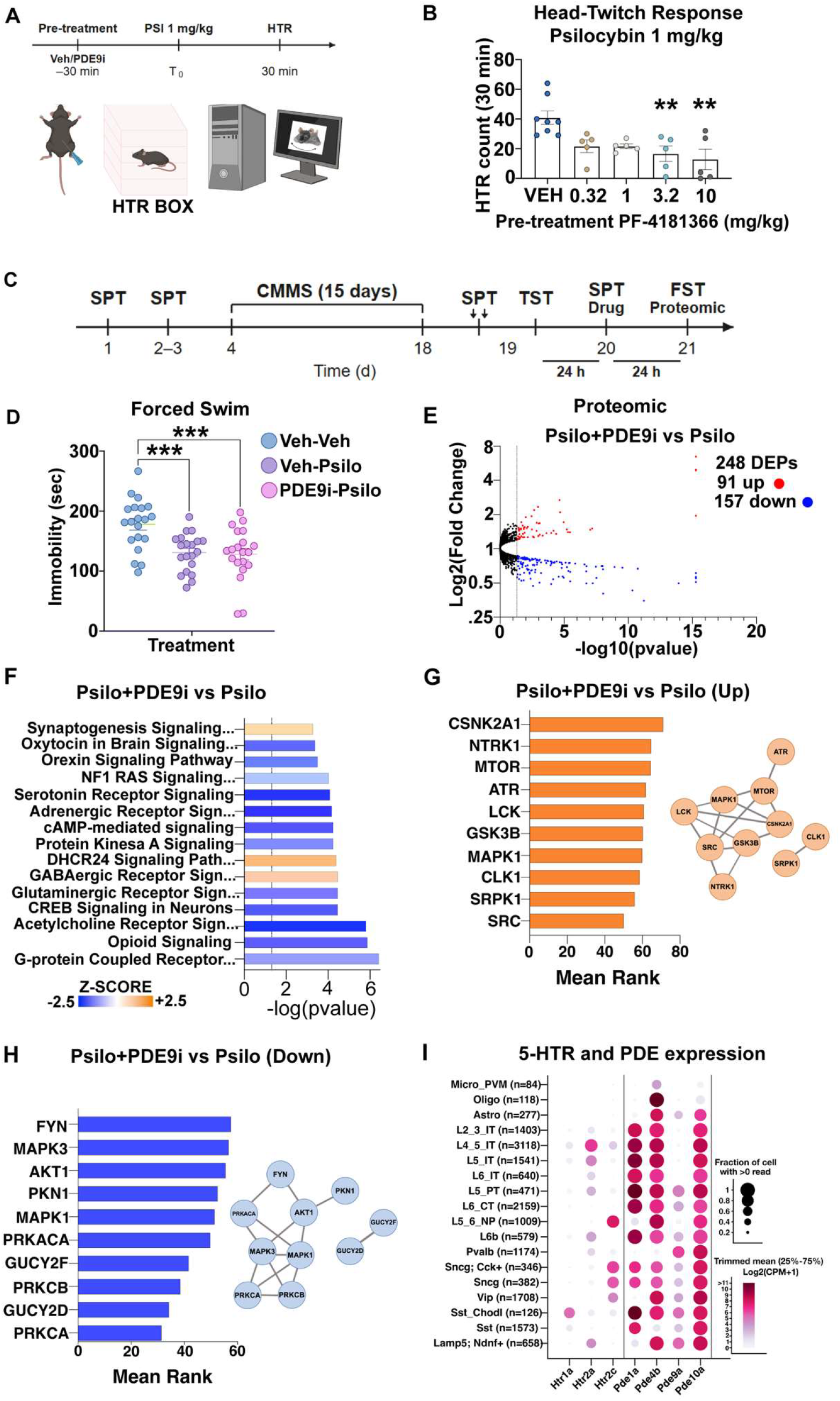
PDE9 inhibition attenuates psilocybin-induced head-twitch responses without affecting antidepressant efficacy, and preserves synaptogenic signaling pathways. (A) Experimental timeline for head-twitch response (HTR) assessment. Mice received PF-4181366 or vehicle 30 minutes prior to psilocybin (1 mg/kg, i.p.) and HTR was recorded for 30 minutes in a dedicated HTR box. (B) Total HTR counts over 30 minutes showing dose-dependent attenuation following PF-4181366 pretreatment. Significant reductions were observed at 3.2 and 10 mg/kg compared to vehicle (**p<0.01; n=5–8/group). (C) Experimental timeline for chronic multimodal stress (CMMS) and behavioral testing. Mice underwent SPT training (day 1), baseline SPT (days 2–3), 15 days of CMMS (days 4–18) with SPT on days 17–18, followed by TST (day 19), drug treatment with SPT (day 20), and FST (day 21; n=20/group). A separate cohort was used for synaptosomal proteomic analysis (n=6/group). (D) FST immobility time. Both psilocybin alone (Veh-Psilo) and in combination with PDE9i (PDE9i-Psilo) significantly reduced immobility compared to vehicle-treated stressed controls (***p<0.001). (E) Volcano plot of differentially expressed synaptosomal proteins (DEPs) in the Psilo+PDE9i group versus Psilo alone (248 DEPs total; 91 upregulated, red; 157 downregulated, blue; n=6/group). (F) Ingenuity Pathway Analysis (IPA) of the Psilo+PDE9i vs. Psilo comparison showing significantly regulated canonical pathways ranked by −log(p-value). Bar color reflects predicted activation (orange) or inhibition (blue) based on z-score (scale: −2.5 to +2.5). (G) Kinase Enrichment Analysis (KEA3) of upregulated DEPs in the Psilo+PDE9i vs. Psilo comparison. Inset: STRING network depicting known protein–protein interactions among predicted upstream kinases, including MTOR, MAPK1, ATR, LCK, SRC, GSK3B, CLK1, SRPK1, NTRK1, and CSNK2A1. (H) KEA3 analysis of downregulated DEPs in the Psilo+PDE9i vs. Psilo comparison. Inset: STRING network of interactions among predicted upstream kinases, including AKT1, FYN, PKN1, MAPK1, MAPK3, PRKCA, PRKCB, PRKACA, GUCY2F, and GUCY2D. (I) Dot plot showing expression of serotonin receptor genes (Htr1a, Htr2a, Htr2c) and phosphodiesterase isoforms (Pde9a, Pde1a, Pde4b, Pde10a) across cortical cell classes and subtypes in the mouse cortex (Allen Brain Cell Atlas). Dot size indicates fraction of expressing cells; color reflects trimmed mean Log2(CPM+1) expression intensity. Data are presented as mean ± SEM. HTR data were analyzed by one-way ANOVA followed by Tukey’s multiple comparisons test. Forced swim data were analyzed by one-way ANOVA with Dunnett’s post hoc test.

## Author Contributions

G.F. and S.J.J. contributed equally to this work. G.F. and S.J.J. designed and performed experiments, analyzed data, and wrote the manuscript. J.R. performed experiments. F.S.M. provided the PDE9 inhibitor and contributed to study design. A.C.K. contributed single-cell transcriptomic analyses and edited the manuscript. J.P.D.A. and J.H.K. contributed to data interpretation and edited the manuscript. C.P. and A.P.K. jointly supervised the work, conceived the study, secured funding, and edited the manuscript. All authors approved the final version and agree to be accountable for all aspects of the work.

## Data Availability

The datasets generated and analyzed during the current study are available from the corresponding author upon reasonable request.

